# Fluid flow structures gut microbiota biofilm communities by distributing public goods

**DOI:** 10.1101/2022.11.11.516095

**Authors:** Jeremy P. H. Wong, Michaela Fischer-Stettler, Samuel C. Zeeman, Tom J. Battin, Alexandre Persat

## Abstract

Bacterial gut commensals experience a biologically and physically complex mucosal environment. While many chemical factors mediate the composition and structure of these microbial communities, less is known about the role of mechanics. Here, we demonstrate that fluid flow impacts the spatial organization and composition of gut biofilm communities by shaping how different species interact metabolically. We first demonstrate that a model community composed of *Bacteroides thetaiotaomicron* (*Bt*) and *Bacteroides fragilis* (*Bf*), two representative human commensals, can form robust biofilms in flow. We identified dextran as a polysaccharide readily metabolized by *Bt* but not *Bf*, but whose fermentation generates a public good enabling *Bf* growth. We demonstrate that in flow, *Bt* biofilms share dextran metabolic by-products, promoting *Bf* biofilm formation. By transporting this public good, flow structures the spatial organization of the community, positioning the *Bf* population downstream from *Bt*. We show that sufficiently strong flows abolish *Bf* biofilm formation by limiting the effective public good concentration at the surface. Physical factors such as flow may therefore contribute to the composition of intestinal microbial communities, potentially impacting host health.

## Introduction

Bacterial commensals colonize the physically-complex environment of the gastrointestinal (GI) tract as dense mixed communities. While most species tend to compete for resources and space, they can also engage in positive interactions that promote diversity. Many bacteria secrete compounds such as nutrient chelators^1^, digestive enzymes^2^, and signalling molecules^3^ that promote the fitness of neighboring cells, either from the same or another taxon. The spatial arrangements of different species within biofilms influences how these competitive or cooperative interactions are enforced^4^, ultimately shaping the composition and function of the bacterial community^5^. Spatial structure of communities is however often overlooked in investigations of microbiota, both *in vivo* and *in vitro*.

Interspecies interactions involving public goods are often studied in liquid cultures^6^, where cells, metabolites and secreted molecules are well-mixed. In the GI tract, commensals however grow as spatially-structured, dense communities exposed to fluid flow^7^. In biofilms, flow exerts hydrodynamic forces that impact the positioning of single cells within the community and transport soluble molecules^8,9^. Advection shapes the distribution of molecules secreted by surface-associated bacteria. For example, flow transports signaling molecules, ultimately structuring gene expression in an isogenic community^10^. By carrying metabolic by-products, flow can also influence the relative spatial arrangement of producers and cheaters^11^. Intestinal-like flows impact the composition of a cross-feeding community growing in microchannels^12^. We therefore postulate that hydrodynamic forces shape community structure in the GI tract, thereby influencing microbiota function. Despite *in vivo* observations and simulations, how physiological and mechanical factors together mediate the composition and spatial structure of gut microbiota communities are not entirely understood.

*Bacteroides spp*. are among the most abundant commensals found in the human bowel^13^. *Bacteroides thetaiotaomicron* (*Bt*) is beneficial to human hosts: it confers colonization resistance and produces metabolically-accessible short-chain and organic acids^14^. *Bacteroides fragilis* (*Bf*) also commonly colonizes the human large intestine, conferring health benefits, but can also opportunistically turn pathogenic^15^. *Bf* and *Bt* leverage polysaccharide utilization loci (PULs) to metabolize a wide range of complex glycans. PUL-encoded proteins process complex polysaccharides into metabolically-accessible sugars^16–20^. The genomes of *Bacteroidetes* species frequently encode hundreds of PULs^21^, allowing these commensals to grow on a broad range of dietary or host-derived glycans^16,17,19,22^.

PULs are often composed of a carbohydrate sensor/transcriptional regulator, various surface glycan-binding proteins, a TonB-dependent transporter, and carbohydrate active enzymes such as glycosylhydrolases (GHs), polysaccharide lyases and carbohydrate esterases^23^. Complex fibres often bind to the cell surface where surface-associated GHs then initiate cleavage of the bulky polysaccharides. Resulting oligosaccharides can subsequently enter the cell^18^, but are also released in the extracellular environment as metabolic by-products^23,24^, thereby constituting a public good accessible to the rest of the community. Thus, a species otherwise unable to utilize the original polysaccharide could grow by utilizing this public good^25^. Consistent with this scenario, *Bacteroides* species share metabolic by-product glycans with other species in liquid cultures^25,26^. In the context of the GI tract, flow could transport these metabolites, affecting their distribution and access to other commensals. These changes would impact the spatial growth pattern of secondary degraders.

We hypothesize that flow impacts the spatial organization of syntrophic microbiota communities by transporting metabolic by-product public goods. To demonstrate this, we investigated nutrient sharing between *Bt* and *Bf* biofilms. We show that both species can form robust surface-associated biofilms in flow environments. We found that when degrading dextran, a common food additive^27^, *Bt* sustains *Bf* growth in co-culture by sharing a metabolic by-product. We found that fluid flow strongly impacts the spatial organization and composition of these syntrophic *Bacteroides* biofilm communities.

## Results

### *B. theta* and *B. fragilis* form biofilms in flow

In the intestine, commensals form dense and organized communities associated with the mucosal surface^28^. *Bt* isolates have variable abilities to form biofilm-like macroscale multicellular structures^29^. Whether these multicellular structures are sufficiently cohesive and resistant to flow is unclear^30^. To test *Bt’s* ability to form biofilms in flow conditions that replicate the fluidic environment of the intestine, we implemented a microfluidic-based anaerobic biofilm assay for single-cell level imaging. We initially seeded *Bt* constitutively expressing sfGFP in microchannels, flushed unattached cells and initiated growth under constant flow of Bacteroides minimal medium (BMM) supplemented with glucose for 4 days (Fig. 1A). To visualize colonization patterns, we then imaged the surface-associated population using spinning disc confocal microscopy.

**Figure 1:**
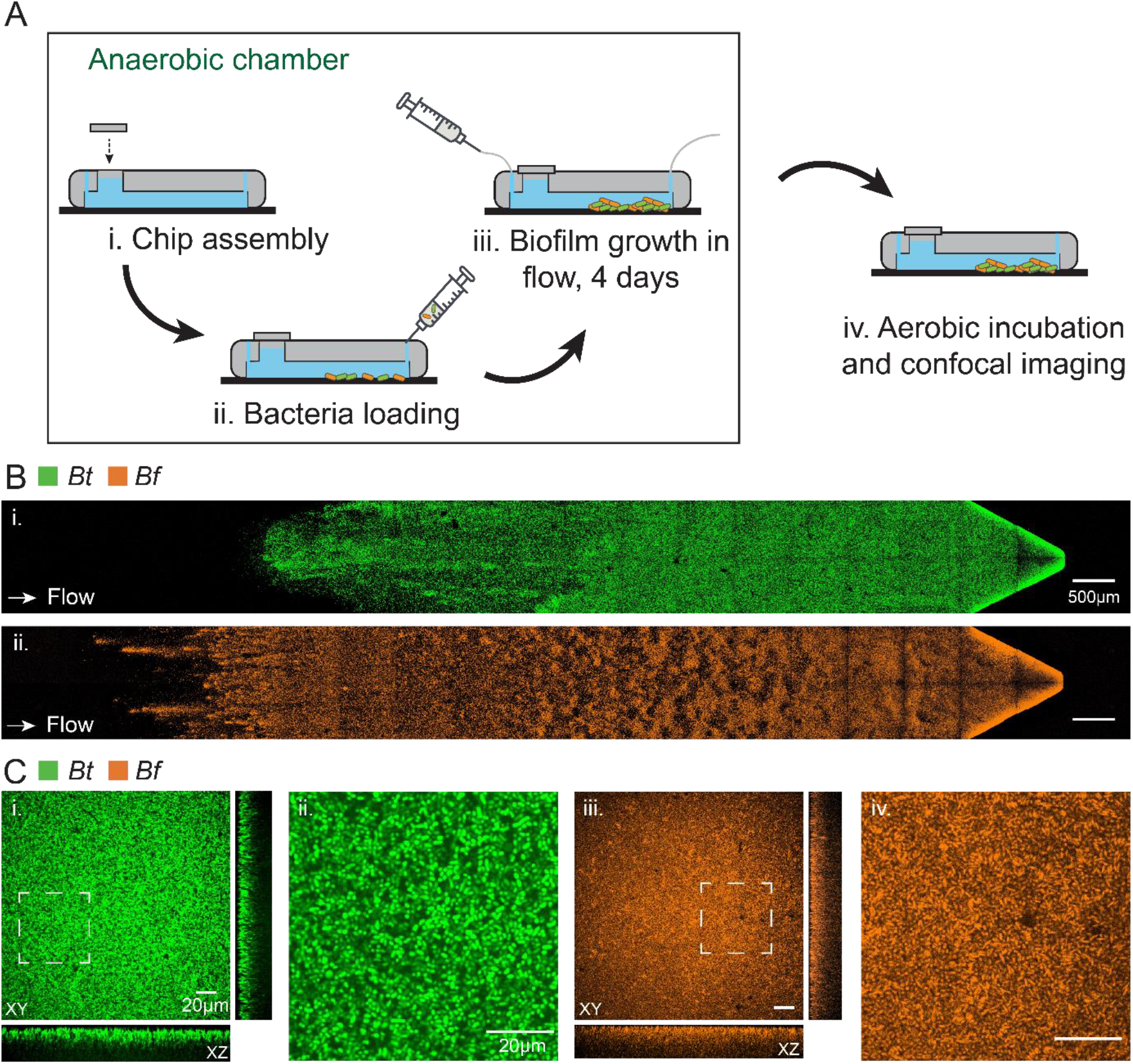
*Bacteroides thetaiotaomicron* (*Bt*) and *Bacteroides fragilis* (*Bf*) form biofilms in flow. (A) Schematic of experimental procedure for anaerobic biofilm growth in flow. (B) Full-channel view of biofilm from (i) *Bt* constitutively expressing sfGFP and (ii) *Bf* constitutively expressing mCherry. (C) Three-dimensional and close-up view of (i, ii) *Bt* and (iii, iv) *Bf* biofilms show the tight packing of single cells and growth in the channel depth. All biofilms were grown anaerobically for 4 days under flow of *Bacteroides* Minimal Medium (BMM) supplemented with glucose.

*Bt* formed cohesive multicellular structures that remained attached to the surface despite flow (Fig. 1B). Visualizations at the single-cell level showed that bacterial cells were contiguous (Fig. 1C). These multicellular structures extended in in the depth of the channel, roughly 10-15 μm above the coverslip surface. These observations show that *Bt* can form surface-attached biofilms that resist shear forces generated by flow (Fig. 1B, C). *Bf* constitutively expressing mCherry also formed dense surface-associated communities of contiguous cells with an architecture resembling the one of *Bt* (Fig. 1B, C). Our results suggest that multiple members of the *Bacteroidota* phylum can colonize the GI tract in the form of mucosa-associated biofilms. We subsequently hypothesize that flow-induced transport near biofilms modulates the concentration landscape of nutrients and ultimately growth. To test this hypothesis, we investigated how a biofilm community composed of *Bt* and *Bf* self-organizes when sharing or competing for nutrients in flow.

### Cross-feeding between B. theta and B. fragilis

*Bacteroides* species possess dedicated PULs that permits growth on a wide-range of specific polysaccharides^31^. *Bt* and *Bf* are versatile utilizers with 84 and 57 predicted PULs, respectively^32^. Some of these systems process the same polysaccharides in *Bt* and *Bf*, so that *Bt* and *Bf* can compete for a single dietary fiber. Others are specific, in principle promoting the growth of only one of the two species. However, polysaccharide degradation can release metabolic by-products accessible to the second species. We therefore searched for nutrient-sharing conditions by identifying a polysaccharide that *Bt* or *Bf* alone can metabolize to generate a by-product that can sustain growth of the other. Based on previous measurements of *Bacteroides* growth under various nutrient conditions, we selected three candidate polysaccharides susceptible to induce nutrient-sharing: dextran, arabinan and inulin^17^,^33–35^.

We assessed the growth of *Bt* and *Bf* in test tubes as mono- and co-cultures in BMM media supplemented with each polysaccharide. *Bt* efficiently grew on dextran and arabinan, while *Bf* could not (Fig. 2A). Conversely, *Bf* grew on inulin, but not *Bt*. These nutrients therefore promote the exclusive growth of a single species out of the two, illustrating specificity in polysaccharide utilization by *Bt* and *Bf*. To investigate whether utilization of one of the three candidate glycans could stimulate nutrient sharing, we co-cultured *Bt* and *Bf* in BMM supplemented with either inulin, arabinan or dextran. We mixed *Bf-*mCherry and *Bt-* sfGFP at a 1:1 ratio in the initial inoculum. To identify nutrient sharing conditions, we quantified the relative population of *Bt* and *Bf* in these cultures using fluorescence microscopy. After 1 and 2 days of growth in anaerobic conditions, we sampled the cultures, incubated them at atmospheric conditions, imaged to count mCherry- and sfGFP-positive cells, and finally computed the *Bf* fraction. In competition for glucose, *Bt* and *Bf* grew roughly in balance, with only a slight advantage for *Bf*. In arabinan, the fraction of *Bf* dropped after 1 day, showing it is likely unable to utilize *Bt’s* arabinan metabolic by-products (Fig. 2B). Conversely, *Bf* showed a strong growth advantage over *Bt* in inulin, suggesting that *Bf* does not produce glycans accessible to *Bt*. The consortium showed a striking difference in dextran: the fraction of each species in co-culture remained close to 50% throughout the 2 days of growth. Growth was balanced in dextran of low and high molecular weight alike, suggesting that *Bt’s* capability of utilizing dextran is independent of the polymer’s structure^36^. These results suggest that, while it is unable to grow alone, *Bf* can benefit from *Bt’s* growth in dextran during co-culture. Consistent with this scenario, *Bf* grows in *Bt* spent dextran medium (Fig. S1).

**Figure 2:**
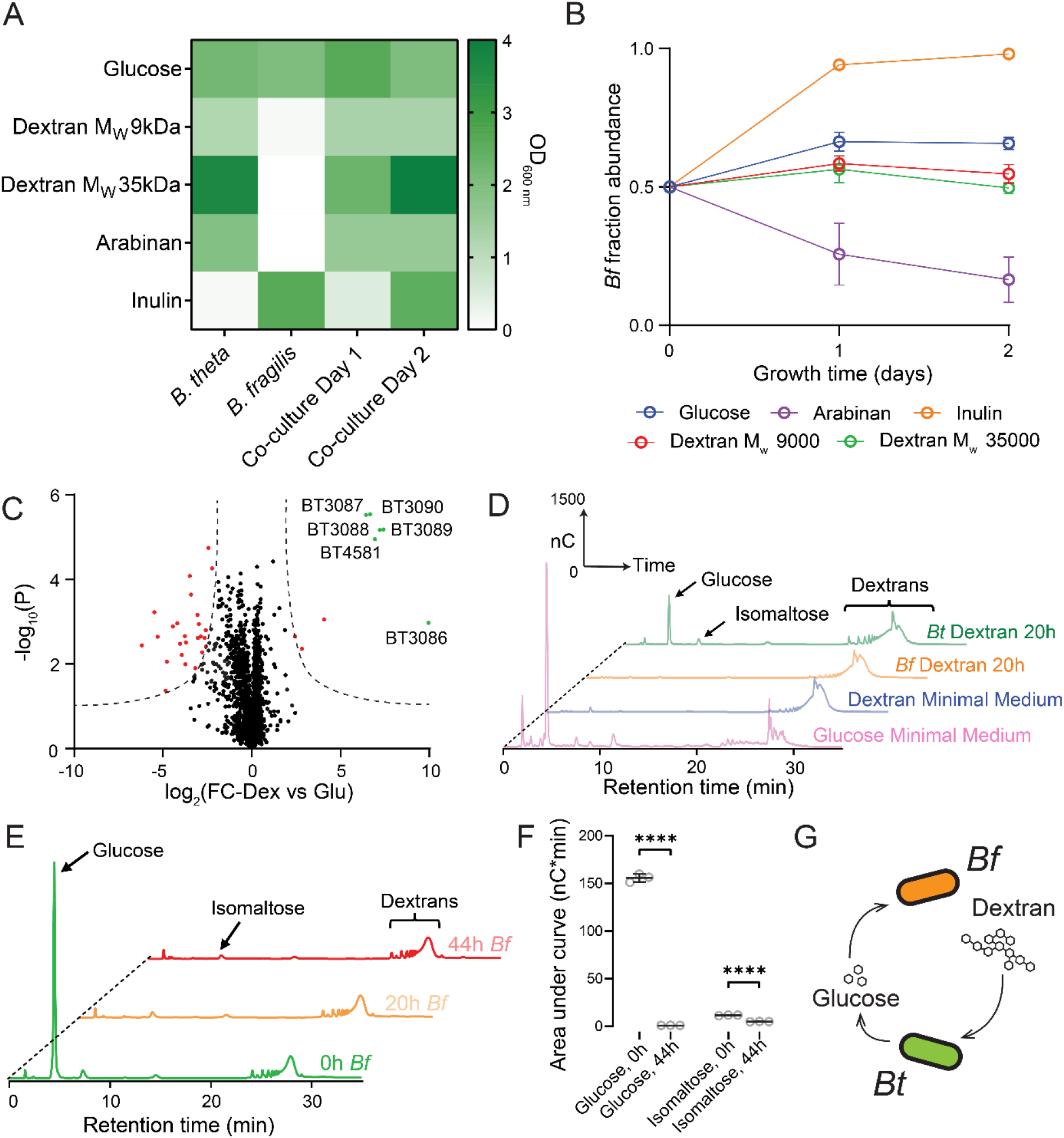
*Bt* shares a dextran degradation by-product that enables *Bf* growth in co-cultures. (A) *Bt, Bf* growth in mono- and co-cultures in dextran, arabinan and inulin. We measured optical density (OD) after 2 days. (B) Fraction of *Bf* in co-culture with *Bt* in the different glycan sources measured by fluorescence imaging (*N* = 3, error bars: standard deviation). (C) Proteomic analysis of *Bt* growing in dextran. We generated the volcano plot based on fold-change in dextran relative to culture in glucose (*N* = 3). Red and green dots correspond to proteins with statistically significant different fold changes (S0 value of 1.0 and FDR cut-off of 0.01). Proteins from genes encoding for PUL48 are strongly upregulated (green dots). (D) HPAEC-PAD analysis of the metabolic output of *Bt* and *Bf* liquid monocultures in dextran. *Bt* growth in dextran produces metabolites whose retention times are identical to glucose and isomaltose. We could not detect any dextran degradation by *Bf*. (E) HPAEC-PAD analysis of the metabolic output of *Bf* liquid monoculture in *Bt* spent medium. The amplitude of the glucose peak decreases during *Bf* growth. (F) Area of the glucose and isomaltose peaks during *Bf* growth in spent medium. (*N* = 3, error bars: standard deviation). (G) Schematic of nutrient sharing from *Bt* to *Bf* in dextran. *Bt* metabolizes the polysaccharide dextran and releases glucose, which is cross-fed to *Bf*. (D-F) HPAEC-PAD experiments were run in triplicates, we display one representative chromatogram.

To verify that *Bf* grows through exploitation of a public good, we investigated *Bt’s* physiology during dextran fermentation. We first compared the proteomes of *Bt* growing in dextran and glucose. Differential expression analysis shows that *Bt* strongly upregulates a group of proteins in dextran (Fig. 2C). These proteins are encoded in an operon (BT3086-BT3090) predicted to belong to the PUL48 of *Bt*^32^. These genes encode for homologues to the nutrient binding protein SusD, the importer protein SusC, a SusE homologue and 2 GHs: α-glucosidase II and dextranase. This operon had previously been identified as essential for *Bt’s* growth in dextran in a transposon screen^37^. Alongside PUL48, we also identified a sole GH, BT4581, encoding for an α-glucosidase which is also highly upregulated in dextran growth. Dextran is a polyglucan with α-1,6 and α-1,3 glycosidic linkages. Since the dextranase BT3087 and α-glucosidases are known to liberate glucose as a product^38^,^39^, we therefore hypothesized that *Bt* degrades dextran into glucose or small gluco-oligosaccharides which promote *Bf* growth.

To expose the nature of nutrient sharing, we analyzed the metabolites released by *Bt* during dextran metabolism. We performed High-Performance Anion Exchange Chromatography - Pulsed Amperometric Detection (HPAEC-PAD) analysis of BMM-dextran growth media. Chromatograms of sterile BMM supplemented with glucose and dextran showed dense peaks at short and long retention times, respectively. The chromatograms of supernatant from *Bt* cultures in BMM-dextran showed an increased density of peaks at short retention times compared to the negative control (Fig. 2D). The most prevalent peak induced by *Bt* growth matched the retention time of glucose. To test whether *Bf* utilizes glucose produced by *Bt* as public good, we inoculated *Bf* into *Bt* spent medium and analyzed the supernatant after 20 h and 44 h of incubation (Fig. 2E). HPAEC-PAD analysis showed that the glucose peak decreases in amplitude after *Bf* growth. We estimated glucose utilization from spent medium by integrating the glucose peak area. Peak area decreased sharply as a function of time of *Bf* incubation, in comparison to the isomaltose peak (Fig. 2E, F). Our results show that when metabolizing dextran, *Bt* releases glucose which cross-feeds *Bf* (Fig. 2F). Therefore, *Bf-Bt* co-culture in dextran therefore represents a realistic model for investigating the feedback between spatial structure and flow during public good sharing between microbiota biofilms.

### Nutrient sharing structures Bacteroides biofilms

In any fluidic environment, molecules secreted by biofilm-dwelling bacteria diffuse and go with the flow. The relative contribution of these two transport mechanisms impacts distribution and ultimately their effective concentration near biofilms. We therefore compared the architectures of *Bt-Bf* biofilm communities while sharing public goods in the absence and presence of flow. We observed that *Bt* formed biofilms under flow of dextran with similar density and architecture as in glucose (Fig. 3A). In the same conditions, *Bf* alone attached to the channel surface but failed to grow into biofilms, as expected from growth in test tubes (Fig. 3B). By contrast, *Bf* formed biofilms when co-cultured with *Bt* (Fig. 3C). While *Bt* uniformly colonized the surface in both dextran and glucose, the spatial arrangement of *Bt* and *Bf* cells in dextran-grown biofilms was markedly different from conditions without cross-feeding (Fig. 3C, D). In particular, the nearest neighbour distance between *Bt* and *Bf* is higher in dextran than glucose (Fig. 3C). To highlight the effect of nutrient-sharing on *Bt* fitness, we measured how *Bf* influences *Bt* biofilm growth. We compared *Bt* biomass accumulation in the presence and absence of *Bf*. We could however not distinguish any difference in *Bt* colonization between those two conditions (Fig. S2).

**Figure 3:**
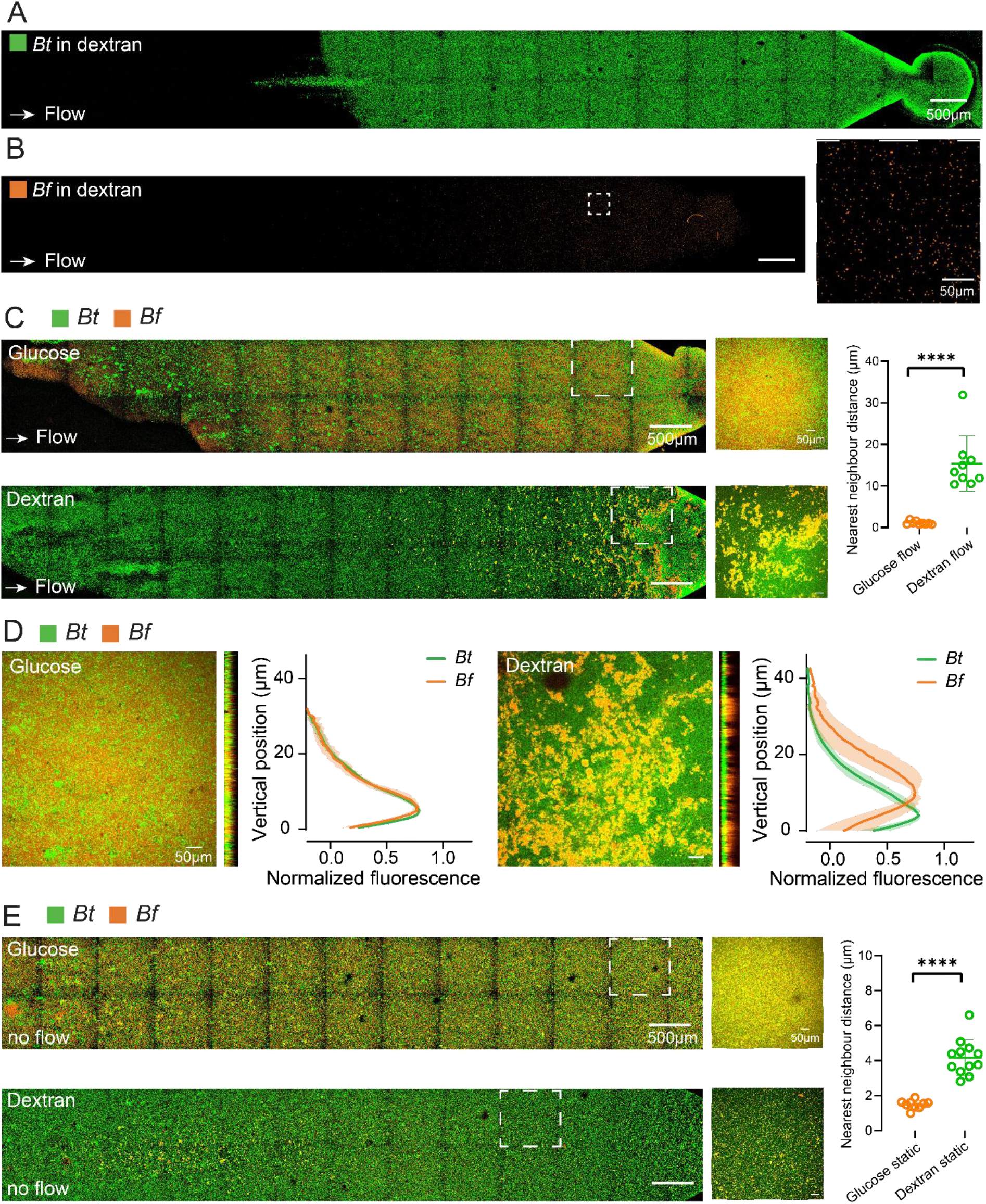
Flow structures syntrophic *Bt-Bf* biofilm communities at the microscale. (A) *Bt* biofilm growing in dextran. (B) *Bf* does not form biofilms in dextran. However, individual cells still remain attached to the glass surface during the 4 days of incubation in flow (close-up view on the right). (C) 4-day *Bt-Bf* mixed biofilms in flow of glucose (top) and dextran (bottom). Nearest neighbour analysis of co-culture growth in dextran and glucose show increased segregation during cross-feeding. (D) Three-dimensional structure of 4-day coculture biofilm in glucose and dextran in flow. We computed the fluorescence profile of the biofilm as a function of height. In glucose, *Bt* and *Bf* grow at the coverslip surface, while *Bf* grows on top of *Bt* biofilms in dextran (*N* = 3, shaded regions: standard deviation across replicates). (E) 4-day *Bt-Bf* mixed biofilms of in glucose and dextran without flow. Nearest neighbour analysis of coculture growth in dextran and glucose again show increased segregation during cross-feeding. However, the degree of segregation difference between the two conditions is less compared to in the presence of flow. All images are representative (*N* = 9). See figure 4 for quantification and statistical significance.

*Bf* microcolonies only noticeably grew after 3 days of co-culture while *Bt* already formed biofilms early on (Fig. S3). This suggests that the *Bt* population density must reach a critical value to generate sufficiently high concentration of metabolic by-products to enable *Bf* growth. Consistent with this sequence of events, *Bf* biofilms mainly grew on top of *Bt* biofilms, whereas in glucose they tend to grow on the glass surface (Fig. 3D). In summary, sharing nutrients imposes spatial and temporal constraints on biofilm growth in the presence of flow. To reveal the contributions of flow in shaping the structure of *Bt-Bf* communities, we imaged the same biofilms grown in the absence of flow. The two-species biofilms also formed, but with a more mixed architecture all the way down to the single cell level. Again, the nearest 1^st^ neighbour distance between *Bt* and *Bf* is higher in dextran than glucose, but the difference between the two conditions is lower than in flow conditions (Fig. 3C, E). We conclude that flow plays a crucial role in shaping the architecture of the *Bacteroidetes* biofilm communities. We therefore suspect that flow mediates the organization and composition of this community by distributing public goods at larger scales.

### Flow shapes community composition

Physiologically, gut contents enter the proximal colon at an estimated high rate of 30 μm/s^40^, where this flow velocity in the colon then drops to about 5 μm/s by the end of the ascending colon^41^. Therefore, to study the impact of fluid flow on microbiota communities in the context of colonic physiology, we grea mixed biofilms a two physiological flow magnitudes: 1.0 μL/min (strong flow) and 0.1 μL/min (weak flow), corresponding to average flow velocities of approximately 28 μm/s and 2.8 μm/s respectively.

While *Bf* and *Bt* uniformly colonized the surface in resting fluid, a weak flow shifted *Bf* colonization downstream of *Bt* (Fig. 4A, B). This observation is consistent with a mechanism where flow transports *Bt-* produced public goods. We quantified the downstream population shift of *Bf* by defining λ as the distance between growth fronts of *Bt* and *Bf* biofilms (Fig. 4C). In the absence of flow, we found no difference between the population shift in glucose and dextran. By contrast, λ increased under flow of dextran, while it had a mild opposite effect in flow of glucose (Fig. 4D). Our results show that fluid flow transports public goods downstream as they are produced upstream. This allows a secondary utilizer to grow, but in a niche separate from the biofilm of the primary utilizer. Without flow, dextran utilization by *Bt* produces free glucose that diffuses around biofilms, slowly accumulating uniformly to promote isotropic growth of *Bf*. Flow however transports *Bt*-produced glucose downstream. As a result, the effective concentration of secreted glucose increases along the channel^10^. Only when *Bt* biofilms reach a sufficient density will they produce an amount of glucose promoting *Bf* biofilm growth.

**Figure 4:**
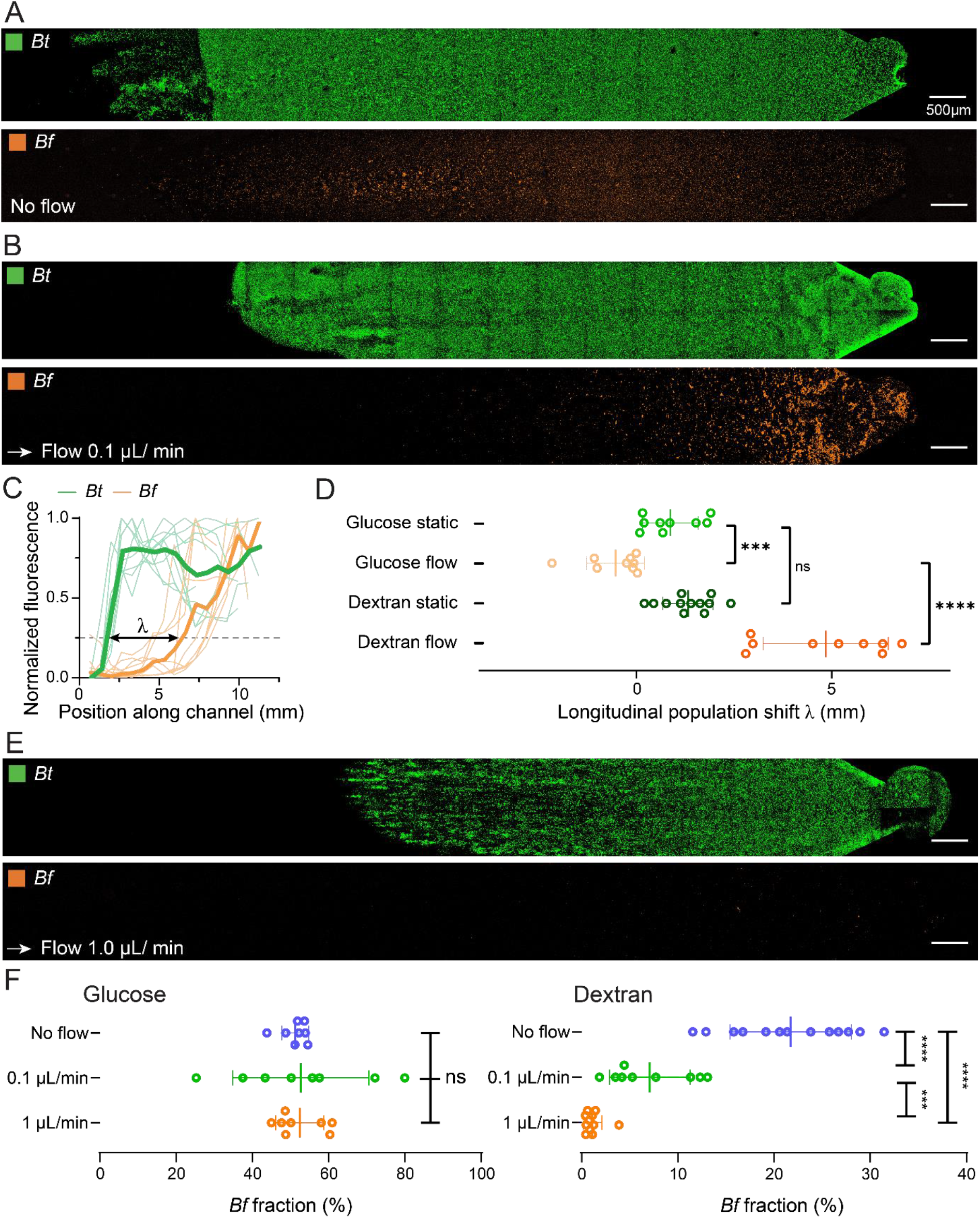
Flow impacts biofilm community composition and organization by transporting public goods. *Bf* and *Bt* biofilms in co-culture grown in dextran for 4 days (A) without flow and (B) with flow (flow rate = 0.1 μL/min). (C) Cross-sectional averaged fluorescence intensity profiles of *Bf* and *Bt* biofilms along the microchannel. Each line corresponds to a biofilm from a single co-culture experiment. Bold lines correspond to the average of 9 replicates. We define the longitudinal population shift *λ* as the *x*-position difference between the *Bt* and *Bf* population across the channel at 25% of the maximal normalized fluorescence intensity. (D) *Bf* longitudinal shift (*λ*) of co-culture biofilms grown in dextran or glucose, in the presence and absence of flow. (E) Confocal images of *Bt* and *Bf* co-culture biofilms grown in dextran for 4 days in strong flow (flow rate, 1 μL/min). (F) Fraction of *Bf* in co-culture biofilms grown in glucose or dextran at different flow rates. *Bf* fraction decreases with flow in dextran cross-feeding conditions, while it remains insensitive to flow in glucose. Error bars show standard deviation across replicates. Statistical test: unpaired t-tests, *** 0.001 < *p* < 0.0001 and **** *p* < 0.0001.

We hypothesized that high flow conditions not only transport but even deplete nutrients from the local environment so that the secondary utilizer cannot grow anymore. To test this hypothesis, we further increased the flow of incoming medium to reduce the effective public good concentration in the microchannel. In strong flow, *Bf* cells remained attached but failed to grow on the surface as biofilms (Fig. 4E). We quantified the effect of flow on the stability of *Bf* by computing the relative abundance of *Bf* and *Bt*. Both species grew equally in glucose irrespective of flow intensity (Fig. 4F). In dextran, the *Bf* fraction was however lower than *Bt*, and decreased as flow increased (Fig. 4F), so that *Bf* was unable to form biofilms at the highest flow rate (Fig. 4E). These results further demonstrate that flow can shape the landscape of public good availability to the point where it can eliminate one of the species from the system.

## Discussion

In well-mixed liquid culture communities, social interactions impact the growth of each member, ultimately guiding ecosystem dynamics. Metabolic interactions are common among microbiota species and provide immediate benefits to the community and the host. For example, commensals share acetate and butyrate, the end products of carbohydrate metabolism^42^. *In vitro, Bacteroides ovatus* produces enzymes which degrades complex polysaccharides into oligosaccharides on the cell surface, which can be utilized by surrounding bacteria^25^. We here found that *Bt* can share dextran metabolic by-products with *Bf*, highlighting an additional syntrophic interaction between gut microbiota species.

In the context of biofilms, population structure mediate social interactions between different clones or species^4^. Surface attached biofilm communities in the GI tract also experience mechanical forces generated by the flow of gut content and mucus, which in principle should impact spatial organization and community composition^41^. Our results show that *Bt* shares public goods with *Bf* within biofilms in a flow-dependent manner. These flows replicate the ones which the microbiota experiences in the proximal and distal colon. At intermediate flow, flow shifts *Bf* biofilms downstream from *Bt* biofilms. In stronger flow, advection reduces the local concentration of metabolic by-product to such an extent that it abolishes *Bf* colonization. Thus, by influencing the transport of signalling molecules or public goods, fluid flow mediates social interactions and ultimately biofilm community composition (Fig. 5)^43^. Our results are therefore consistent with computational and experimental investigations predicting that flow spatially structures microbiota and ultimately affects composition^12,41,44^.

**Figure 5:**
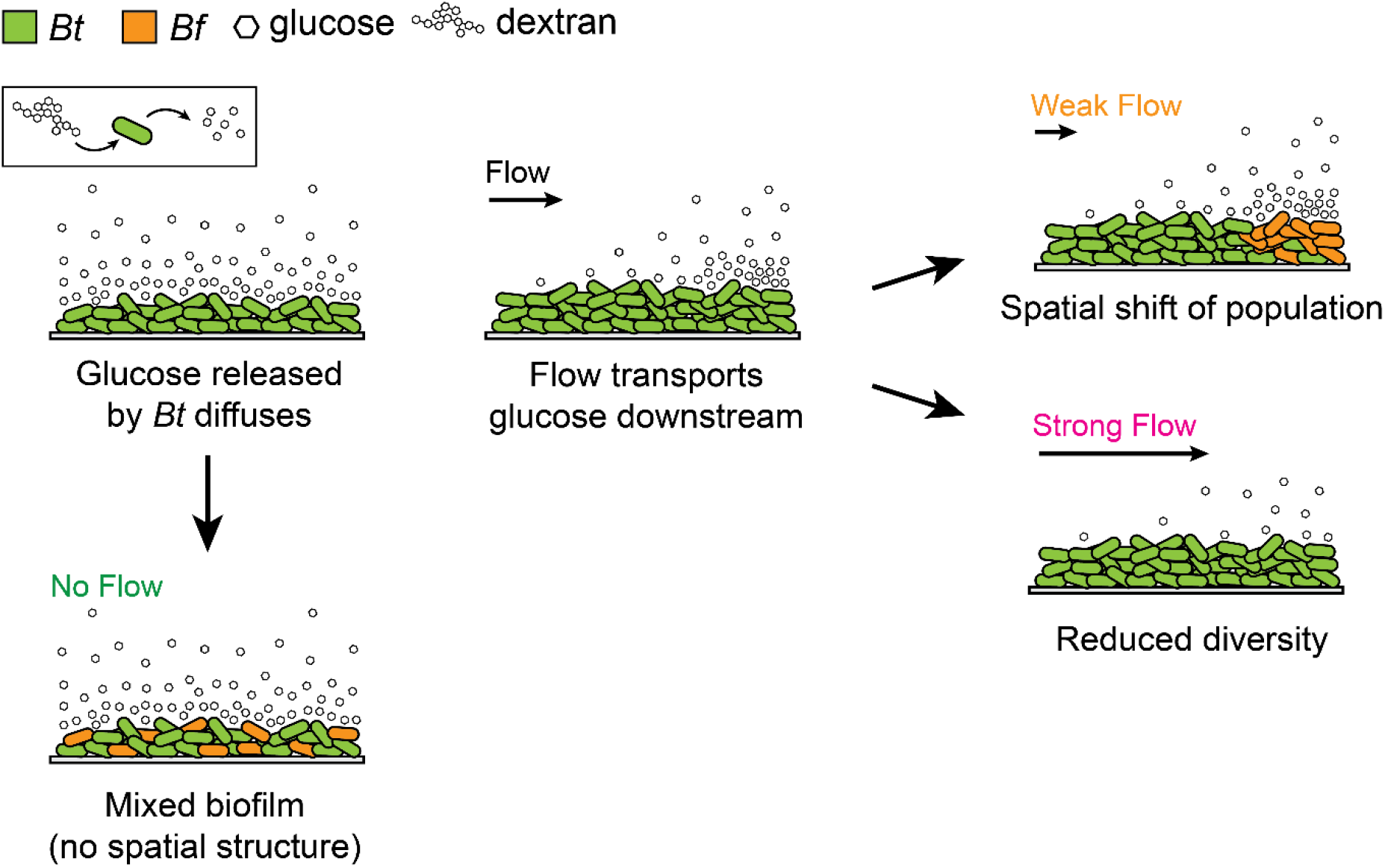
Proposed mechanism of how fluid flow structures nutrient sharing gut microbiota biofilm communities. *Bt* releases glucose when metabolizing dextran. Without flow, glucose diffuses anisotropically. Fluid flow transports glucose downstream, generating a concentration gradient along the channel. *Bf* grows downstream in regions with high glucose concentration. In sufficiently strong flow, glucose concentration is not sufficient to sustain *Bf* growth. Thus, flow impacts community composition by shaping the chemical landscape of public goods.

By controlling the physical and chemical environment of the mucosa, the host creates distinct ecological niches along the GI tract^45^, each of which is colonized by a specific set of species^46^. Thus, gradients of pH, oxygen or mucus production from the stomach to the colon impose local selective pressures that drive the spatial organization of the microbiota. This spatial organization generates chemical gradients of metabolites and signalling molecules that in turn impact host physiology, ultimately shaping mucosal environment. The spatial structure of microbiota communities thus mediates a feedback between host and commensals, ultimately influencing intestinal homeostasis and host physiology.

We postulate that fluid flow mediates the spatial organization of bacteria along the human colon in part by influencing the distribution of carbohydrates. Commensals capable of utilizing complex polysaccharides have a selective advantage in colonizing the proximal colon. Utilizers of simpler glycans produced by primary fermenters would thus colonize the more distal colonic regions. Consistent with this, the overall microbial abundance and diversity increases from the proximal to the distal colon^7^. By transporting metabolites downstream of primary fermenters, flow strongly impacts community composition and can even abolish the growth of a secondary utilizer. Therefore, the biofilm lifestyle and flow are crucial factors to consider in investigations of microbiota composition and stability.

With a realistic model microbiota that integrates polysaccharide utilization and cross-feeding in biofilms, we demonstrated that flows at physiologically-relevant velocities influence the architecture and composition of microbiota biofilms. We therefore postulate that hydrodynamic perturbations induced by changes in intestinal flows or viscosity in the GI tract will impact metabolic interactions between commensals, thereby modulating community structure and host physiology. The complex mechanisms that mediate the spatial organization of more complex microbiota communities are still not entirely identified, but likely combine biochemical and physical factors. Advanced *in vivo* imaging methodologies of microbial communities have the potential to provide a high-resolution perspective on these complex communities and how they assemble at the host mucosal surface^28^,^47^. A mechanistic understanding of these principles will however require combining animal studies with *in vitro* experimentations. The development of host models integrating critical *in vivo* parameters such as epithelia and its mucus layer will ultimately allow us to identify the forces that shape gut microbiota composition and organization^48,49^.

## Materials and Methods

### Strains and culture media

*Bacteroides thetaiotaomicron* VPI-5482 sfGFP and *Bacteroides fragilis* NCTC-9343 mCherry^47^ were used for all experiments in this work. *Bacteroides* minimal medium (BMM) supplemented with the indicated carbon source was used in all experiments. The medium was prepared as previously described^50^. Briefly, a 10x medium stock containing 1 M KH_2_PO_4_, 150 mM NaCl and 85 mM (NH_4_)_2_SO_4_ was first prepared and adjusted to a pH of 7.2. Solutions of 1 mg/mL vitamin K_3_, 0.4 mg/mL FeSO_4_, 0.1 M MgCl_2_, 0.8% (w/v) CaCl_2_, 0.01 mg/mL vitamin B_12_ and 1.9 mM hematin–0.2 M histidine solutions were prepared separately. To prepare 100 mL of BMM,10 mL of the 10x salts were mixed with 0.1 g of L-cysteine, 50 μL of vitamin B_12_ solution, and 100 μL of each vitamin K_3_, FeSO_4_, MgCl_2_, CaCl_2_ and hematin-histidine solution. All carbon sources were added to a final concentration of 5 mg/mL. The media were filter-sterilized using a 0.22 μm filter unit and degassed in the anaerobic chamber overnight. All bacteria were cultured at 37 °C under anaerobic conditions in a vinyl anaerobic chamber (COY) inflated with a gas mix of approximately 5% carbon dioxide, 90% nitrogen and 5% hydrogen (CarbaGas). Bacteria were first streaked on brain heart infusion plates (SigmaAldrich) containing 10% sheep blood (TCS BioSciences). Single colonies were then inoculated in BMM and grown overnight.

### Microchannels fabrication

Microchannels were fabricated by standard soft-lithography as previously described^51^ with slight modifications. Upon cutting out the cured PDMS (Sylgard 184, Dow Corning) channels (2 cm-long, 2 mm-wide channels, 300μm-deep), 1 mm inlet and outlet, along with a 6 mm hole were punched immediately downstream of the inlet. The PDMS was then bound to cleaned glass without plasma treatment. Medium was then loaded through the inlet until the 6 mm hole was filled up, where a piece of PDMS was then used to seal the hole. This hole served as the bubble trap of the microchannel. Medium was then loaded until the channel was filled.

### Biofilm growth

All dextran biofilm growth experiments were performed in BMM-dextran (M_w_ 35000-45000 Da). For glucose co-cultures, overnight monocultures of *Bt* and *Bf* in glucose were mixed at a 1:1 ratio upon OD measurements and appropriate initial dilutions. For dextran co-cultures, *Bt* and *Bf* were mixed at a 1:1 ratio to a final OD of 0.1 and grown as a co-culture overnight. To initiate biofilm formation in flow, cell cultures were diluted to a final OD of 0.01. 4 μL of the diluted bacterial culture was then loaded into the microchannels through the outlet and statically incubated for 15 minutes promoting initial attachment. The inlet was then connected to a 10 mL disposable syringe (BD Plastipak) filled with the medium and mounted onto a syringe pump (KD Scientific), using a 1.09 mm outer diameter polyethylene tube (Instech) and a 27G needle (Instech). All biofilms were then grown anaerobically for 4 days at 37°C under the volume flow rates of 1.0 μL/min or 0.1 μL/min, which corresponds to the flow velocities of roughly 28 μm/s and 2.8 μm/s respectively. These values were chosen as they fall into the physiological range of flow velocities between the proximal and the end of the ascending colon: approximately 30 μm/s^40^ and 5 μm/s^41^. Upon 4 days of growth, the tubings were carefully disconnected. For biofilm growth in the absence of flow, cells were cultured, and seeded as mentioned above. 10 μL of medium was then injected slowly through the inlet to flush away unattached cells. 100 μL droplets of medium was then carefully deposited on top of the inlet and outlet to prevent the channel from drying. The biofilms were then removed from the anaerobic chamber, and incubated in an oxygenated environment to allow for fluorescent protein folding. The biofilms were then imaged by fluorescence confocal microscopy.

### Microscopy

All biofilm imaging were performed using a Nikon Eclipse Ti2-E inverted microscope coupled with a Yokogawa CSU W2 confocal spinning disk unit and equipped with a Prime 95B sCMOS camera (Photometrics). The two objectives used for imaging were the 20x APOChromat water immersion objective with a numerical aperture (N.A.) of 0.95 and the 60x water immersion objective with an N.A. of 1.20. Z-stacks were recorded at a 0.5 μm step-height. Fiji was then used for the display of all images. For the visualization of the full biofilm, frames were taken across the *xy*-plane and stitched together upon background subtraction as described in the image analysis section. All single cell counting experiments in liquid cultures were performed using the Nikon TiE epifluorescence microscope coupled with a Hamamatsu ORCA Flash 4 camera and a 40 x Plan APO NA 0.9 objective. Fiji was then used to display the images.

### Image analysis

To acquire images of biofilms along the entire channel, tile imaging was performed and images were subsequently stitched together using the ‘make montage’ function upon background subtraction in Fiji. The relative abundance of *Bt* and *Bf* in biofilms were calculated by computing the area of coverage by each strain in Fiji. To quantify the 1^st^ nearest neighbor distance, we selected 4000 random pixels in the segmented sfGFP picture for each stitched biofilm images. We then calculated the distance between the selected pixels and their nearest neighbor pixels in the corresponding segmented mCherry picture.

This way we obtained an average 1^st^ nearest neighbor distance for each stitched biofilm images. For each biofilm, the highest 1^st^ nearest neighbour distance value was filtered out and a mean value of the remaining stacks was calculated. To quantify the relative positioning of *Bf* and *Bt*, we computed the fluorescence intensity along the channel for each 660 μm-wide field of view upon background subtraction. The fluorescent intensity of frames corresponding to the same *x* positions were then averaged. This was then used to generate fluorescent intensity profiles for *Bt* and *Bf*. For each, the biofilm edge was computed as the *x-* position at which the fluorescence reached 25% of the maximum intensity. The difference between *Bt* and *Bf* upstream edge positions were then calculated to measure the positional shift between *Bt* and *Bf*, denoted as λ. A similar procedure without averaging the fluorescent intensity of frames corresponding to the same *x*-positions was used to quantify the relative distribution of *Bt* and *Bf* as a function of biofilm height.

### Competition experiments in test tubes

*Bt* and *Bf* were cultured in Bacteroides Minimal Medium (BMM) supplied with 5 mg/mL of D(+)-glucose (Carl Roth), dextran M_W_ 9000-11000 (SigmaAldrich), dextran M_W_ 35000-45000 (SigmaAldrich), arabinan (Megazyme) and inulin from dahlia tubers (SigmaAldrich). To start competition experiments, *Bt* and *Bf* were grown separately in 1 mL of BMM-glucose overnight. The resulting cultures were then pelleted by centrifugation and washed twice with PBS. The pellets were then resuspended in PBS, where the OD was measured. *Bt* and *Bf* were then mixed at a 1:1 ratio to an initial OD of 0.1 in each of the carbon sources tested. The cultures were then grown anaerobically for 2 days. A microscopy approach was then used to quantify the relative abundances of *Bt* and *Bf* in these cultures. To achieve this, the cultures were mixed well and 10 μL of each cell culture was sampled at day 1 and 2. The cultures were then diluted in PBS and incubated at aerobic conditions. The samples were then spotted onto a glass coverslip and covered by a thin agarose pad prior to fluorescence imaging by widefield fluorescence microscopy. For each sample and replicate, 3 image frames across the agar pad were recorded. The relative abundance of sfGFP- and mCherry-expressing cells in each sample were then quantified using the ‘analyze particle’ function in Fiji and averaged across the 3 frames.

### Sample preparation, mass spectrometry and data analysis for proteomics

*Bt* was cultured in BMM-glucose and BMM-dextran (M_w_ 9000-11000) overnight from an initial OD of 0.01. Cells were lysed and lysates containing 20 μg of protein content were collected for further analysis. Cell lysates were digested with trypsin using the Filter-Aided Sample preparation (FASP)^52^ protocol with minor modifications. Proteins were resuspended in 200 μl of 8 M urea; 100 mM Tris-HCl and deposited on top of Microcon-30K devices. Samples were centrifuged at 9391 g at 20°C for 30 min. All subsequent centrifugation steps were performed using the same conditions. An additional 200 μl of 8 M urea 100 mM Tris-HCl was added and devices were centrifuged again. Reduction was performed by adding 100 μl of 10 mM TCEP in 8 M urea, 100 mM Tris-HCl on top of filters followed by 60 min incubation at 37 °C with gentle shaking and protected from light. Reduction solution was removed by centrifugation and filters were washed with 200 μl of 8 M urea, 100 mM Tris-HCl. After removal of washing solution by centrifugation, alkylation was performed by adding 100 μl of 40 mM chloroacetamide in 8 M urea, 100 mM Tris-HCl and incubating the filters at 37 °C for 45 min with gentle shaking and protected from light. The alkylation solution was removed by centrifugation and another washing/centrifugation step with 200 μl of 8 M urea 100 mM Tris-HCl was performed. This last urea buffer washing step was repeated twice followed by three additional washing steps with 100 μl of 5 mM Tris-HCl. Proteolytic digestion was performed overnight at 37 °C by adding 100 μl of Endoproteinase Lys-C and Trypsin Gold in an enzyme/protein ratio of 1:50 (w/w) on top of filters. Resulting peptides were recovered by centrifugation. The devices were then rinsed with 50 μl of 4% trifluoroacetic acid and centrifuged. This step was repeated three times and peptides were finally desalted on SDB-RPS StageTips^53^.

For LC-MS/MS analysis, resuspended peptides were separated by reversed phase chromatography on a Dionex Ultimate 3000 RSLC nano UPLC system connected in-line with an Orbitrap Lumos (Thermo Fisher Scientific, Waltham,USA). A capillary precolumn (Acclaim Pepmap C18, 3 μm-100 Å, 2 cm x 75 μm ID) was used for sample trapping and cleaning. A capillary column (75 μm ID; in-house packed using ReproSil-Pur C18-AQ 1.9 μm silica beads; Dr. Maisch; length 50 cm) was then used for analytical separations at 250 nl/min over 150 min biphasic gradient. Acquisitions were performed through Top Speed Data-Dependent acquisition mode using a 1 second cycle time. First MS scans were acquired at a resolution of 240000 (at 200 m/z) and the most intense parent ions were selected and fragmented by High energy Collision Dissociation (HCD) with a Normalized Collision Energy (NCE) of 30% using an isolation window of 0.7 m/z. Fragmented ion scans were acquired in the ion trap using a fix maximum injection time of 20 ms and selected ions were then excluded for the following 20 s.

Protein identification and label free quantification were performed using MaxQuant 1.6.10.43^54^. The *B. thetaiotaomicron* Uniprot reference proteome database was used for this search. Carbamidomethylation was set as fixed modification, whereas oxidation (M), phosphorylation (S, T, Y) and acetylation (Protein N-term) were considered as variable modifications. A maximum of two missed cleavages were allowed for this search. “Match between runs” was selected. A minimum of 2 peptides were allowed for protein identification and the false discovery rate (FDR) cut-off was set to 0.01 for both peptides and proteins. Label-free quantification and normalisation was performed by Maxquant using the MaxLFQ algorithm, with the standard settings^55^.

In Perseus^56^, reverse proteins, contaminants and proteins only identified by sites were filtered out. Biological replicates were grouped together and protein groups containing a minimum of two LFQ values in at least one group were conserved. Missing values were imputed with random numbers using a gaussian distribution (width = 0.3, down-shift = 1.8). Two-samples t-test was performed to identify the differentially expressed proteins, followed by a permutation-based correction (False Discovery Rate). Significance curves in the volcano plot corresponded to a S0 value of 1.0 and FDR cut-off of 0.01.

### Glycan analysis using HPAEC-PAD

For culture supernatant analysis experiments, *Bt* or *Bf* was inoculated into 1 mL of BMM-dextran M_w_ 9000-11000 to a final OD of 0.01. The cultures were then grown overnight for 20 hours anaerobically. The resulting cultures were pelleted by centrifugation and the supernatant was collected. Cation and anion exchange chromatography with AmberChrom resins (SigmaAldrich) were then used to clean up the samples followed by filtering through a 0.22 μm sterilization filter prior to analysis. The resulting flow-throughs were then analyzed by HPAEC-PAD. For identification of the carbohydrate cross-fed to *Bf*, we produced *Bt* spent medium from a 30 mL culture grown in BMM-dextran for 20 hours. We pelleted the culture, collected the supernatant, and filtered it using a 0.22 μm sterilization filter. We cultured *Bf* overnight in BMM glucose, washed and resuspended them in BMM without any carbon source, and then added this to the spent medium at a final OD of 0.01. After 24 and 40 hours of growth, we pelleted the cultures and cleaned up the samples with the exact same procedures used for the supernatant samples. The sugars were separated on a Dionex CarboPac PA-20 column from Thermo Fisher Scientific with the following eluent: A) 100 mm NaOH; B) 150 mm NaOH and 500 mm sodium acetate at a flow rate of 0.4 mL/min. The gradient was 0 to 15 min, 100% A (monosaccharide elution); 15 to 26 min, gradient to 10% A and 90% B (malto-oligosaccharide elution); 26 to 36 min, 10% A, 90% B (column wash step); and 36 to 46 min step to 100% A (column re-equilibration). Peaks were identified by co-elution with known glucose, isomaltose standards using the Chromeleon software. The HPLC data was then exported and replotted with the graphing software GraphPad Prism 8, where area under the curve of peaks of corresponding samples were calculated.

## Data availability

The macro used for image stitching, nearest neighbour quantifications and the Matlab code used for the quantification of *Bt-Bf*-population shift will be available on GitHub.

## Acknowledgements

We thank Justin Sonnenburg for strains and plasmids, Liz Shepherd for protocols concerning *Bacteroides* culturing and genetic manipulations, Romain Hamelin and Florence Armand at the proteomics core facility at the EPFL for help with the proteomics experiment and data analysis, Audrey Bender for initial works with nutrient sharing biofilms, and Pascal Odermatt for help with image analysis. This work was supported by SNSF Projects grant 310030_204190 and the EPFL iPhD program.

## Author contributions

J.P.H.W., T.J.B and A.P. conceived the study. J.P.H.W and A.P. designed the study. J.P.H.W. performed all experiments. M.F.S. and S.C.Z performed HPAEC-PAD. J.P.H.W performed all image and data analysis. J.P.H.W created the figures. A.P. and T.J.B. supervised the work. J.P.H.W., T.J.B. and A.P. wrote the paper. All authors provided input on the paper.

---

**Figure S1:**
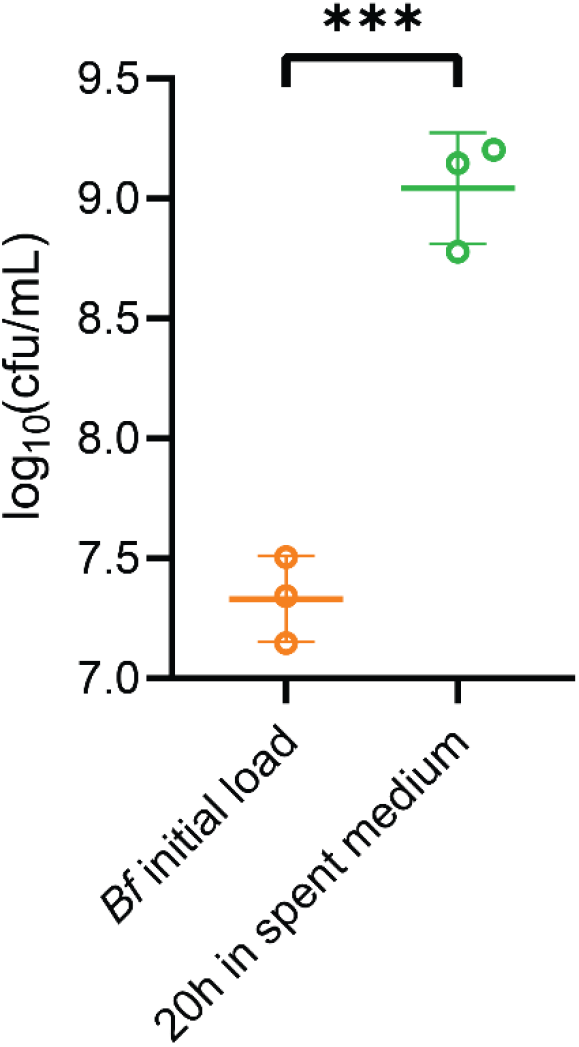
*Bt* spent dextran medium supports the growth of *Bf*. We filter-sterilized a *Bt* monoculture in BMM dextran. We then inoculated *Bf* to this spent medium and performed CFU counts at time of inoculation and after 20 h of incubation. *Bf* density increased more than 10-fold, showing that a public good released by *Bt* through dextran degradation supports *Bf* growth. Error bars show standard deviation across replicates. Statistical test: unpaired t-tests, *** 0.001 < p < 0.0001.

**Figure S2:**
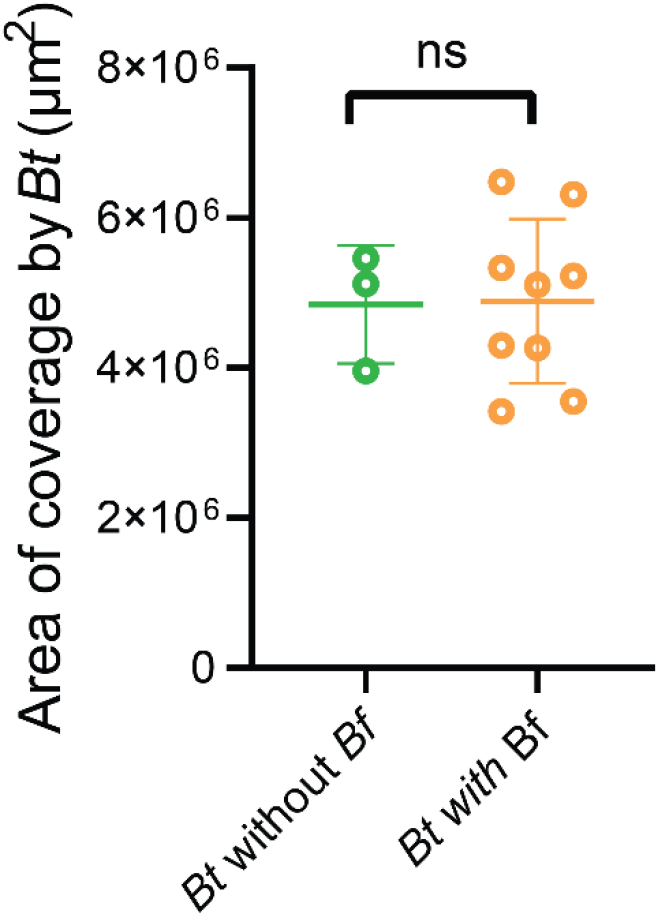
*Bt* biofilm growth in dextran is insensitive to the presence of *Bf*. Area of coverage by *Bt* of monoculture and *Bt-Bf* co-culture biofilms in dextran reveals no significant changes to *Bt* population induced by the presence of *Bf*. Error bars show standard deviation across replicates. Statistical test: unpaired t-test.

**Figure S3:**
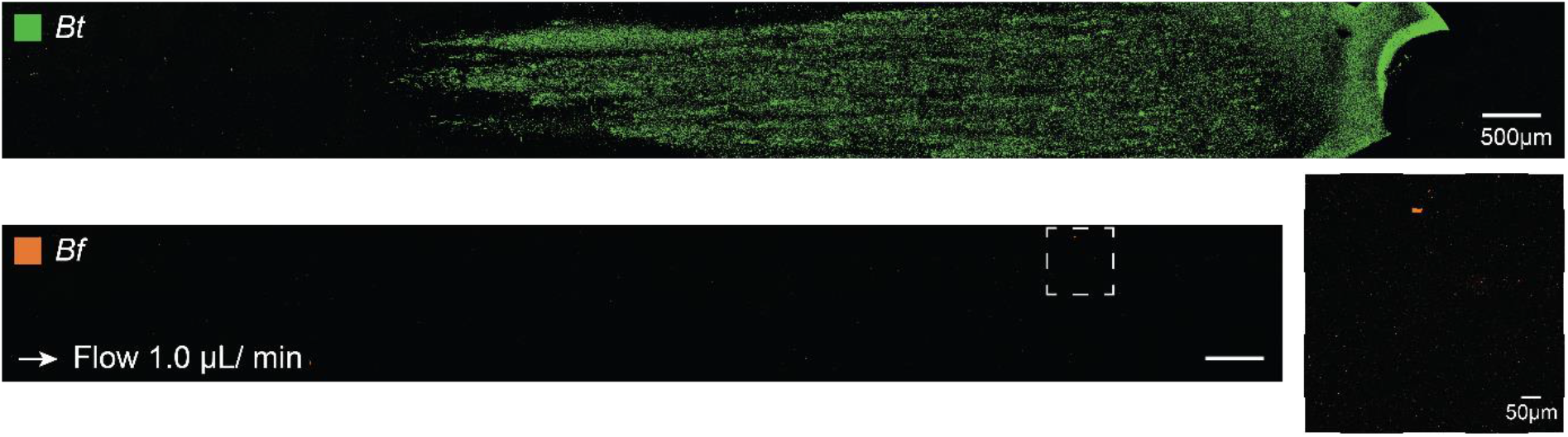
*Bf* growth in co-culture biofilms are delayed compared to *Bt*. After 3 days of co-culture biofilm with *Bt, Bf* shows no measurable growth. This suggests that the density of *Bt* biofilms must reach a critical value to produce sufficiently metabolic by-products to enable *Bf* growth.

